# Ergothioneine and a new food supplement product protect against cyclophosphamide-induced premature ovarian failure in rats

**DOI:** 10.1101/2025.08.30.672979

**Authors:** Chen Jin, Minyan Shi, Han Liu, Siling He, Huawei Zhu, Pan Wang

**Affiliations:** Shenzhen Key Laboratory of Steroid Drug Discovery and Development, School of Medicine, The Chinese University of Hong Kong (Shenzhen); Guangzhou Ruoyuchen Technology Co., Ltd

## Abstract

Premature ovarian failure (POF) is characterized by disrupted estrous cycles and impaired folliculogenesis due to oxidative stress and inflammation. In this study, a cyclophosphamide (CTX)-induced POF rat model was used to evaluate the protective effects of ergothioneine (EGT) and a nutraceutical formula (FineNutri Cellular Vitality Capsules) containing EGT. CTX treatment markedly prolonged the estrous cycle, reduced estrus duration, decreased ovarian weight, impaired follicular development, and increased granulosa cell apoptosis. Serum estradiol (E_2_) and anti-Mullerian hormone (AMH) levels decreased, whereas luteinizing hormone (LH) and follicle-stimulating hormone (FSH) increased, reflecting disruption of the hypothalamic-pituitary-ovarian axis by CTX. CTX treatment induced oxidative stress, with reduced catalase (CAT), superoxide dismutase (SOD) activity, and glutathione (GSH) content, and increased malondialdehyde (MDA) in ovarian tissue. The EGT and the nutraceutical formula restored estrous cycles, increased ovarian weight, improved primordial follicle counts, and reduced atretic follicles and granulosa cell apoptosis. Hormonal balance was partially restored, with increased E_2_ and AMH and reduced LH and FSH levels. Oxidative stress was alleviated with higher CAT, SOD, and GSH levels and reduced MDA concentrations. In addition, EGT and the formula reduced inflammation in skin tissue. These findings suggest that EGT and the nutraceutical formula could protect against CTX-induced POF and help preserve ovarian function, probably by mitigating oxidative stress.

## INTRODUCTION

Premature ovarian failure (POF), also known as primary ovarian insufficiency (POI), is a common gynecological endocrine disorder characterized by amenorrhea, infertility, hypoestrogenism and elevated gonadotropin levels in women under the age of 40 (1). The etiology of POF is multifactorial, involving genetic predispositions, metabolic dysfunctions, pharmacological exposures and autoimmune abnormalities, which are not fully understood yet (2). Current therapeutic strategies for POF are symptomatic, primarily aim to restore, preserve or replace ovarian function, but no effective therapy has yet been established (3). While hormone therapy is the primary treatment for POF with some clinical efficacy, its long-term use may increase the risk of cardiovascular disease, thromboembolism, breast cancer, endometrial carcinoma and other complications (4). Therefore, it is important to discover novel agents for mitigating POF.

Studies have found that oxidative stress is important for POF pathogenesis (5). The most widely accepted theory of ovarian aging is that oxidative stress damages the ovarian granulosa cells and results in a decline in both the quantity and quality of follicles (7,8). In recent years, some studies have found a variety of natural bioactive compounds which have potential in protecting ovarian function. For example, Coenzyme Q10 (CoQ10), a vital coenzyme in the mitochondrial electron transport chain, has potent antioxidant properties and maintains mitochondrial energy metabolism (9, 10). Studies have shown that CoQ10 can improve mitochondrial function in granulosa cells and reduce oxidative stress in oocytes, thereby delaying ovarian aging and improving ovarian reserve (11). In addition, CoQ10 promotes intra-ovarian steroidogenesis, increases follicle number, and improves oocyte quality (12). Pyrroloquinoline quinone (PQQ) is a novel water-soluble redox cofactor with strong free radical scavenging capacity and stimulation of mitochondrial biogenesis (13). It is found that PQQ in combination with mesenchymal stem cells mitochondria ameliorates CTX-induced POF through activation of the SIRT1/ATM/p53 signaling pathway (14). Hydroxytyrosol, one of the most potent phenolic antioxidants found in olive oil, exerts significant free radical scavenging and anti-inflammatory effects (15). Evidence indicates that hydroxytyrosol may inhibit ovarian inflammaging via selective autophagy (16). Furthermore, studies have shown that selenium and vitamin E can support ovarian function recovery in patients with POF (17). Ergothioneine (EGT), a diet-derived sulfur-containing amino acid, has various physiological activities, including free radical scavenging, detoxification, immunomodulation, maintenance of DNA biosynthesis and normal cell growth (18). However, whether EGT has protective effect on ovarian function is not clear.

The study investigates the effects and mechanisms of a composite nutraceutical formulation (FineNutri Cellular Vitality Capsules produced by Finenutri International Brand Management Co., Ltd, Hong Kong, China), hereafter referred to as ‘product’, with each capsule containing 15 mg EGT, 41.5 mg CoQ10, 10 mg PQQ, 5.3 mg hydroxytyrosol and 100 mg vitamin E) and its ingredient EGT on a cyclophosphamide (CTX)-induced POF rat model to determine whether they can protect ovarian function. CTX is a commonly-used inducer of POF (19). These results may contribute to the development of ovarian anti-aging products for ovarian function support and anti-aging nutritional interventions in sub-health women, women recovering ovarian function after chemotherapy, and perimenopausal women.

## MATERIALS AND METHODS

### Animal and treatment

Female SD rats (body weight of 180-210 g) were purchased from Beijing Vital River Laboratory Animal Technology Co., Ltd. and housed in the specific pathogen-free (SPF) condition with a 12:12 hour light/dark cycle and controlled temperature and humidity (22 ± 2 ℃, 50%–60% humidity). All the animal protocols were approved by the Institutional Animal Care and Use Committee (IACUC) of the Chinese University of Hong Kong (Shenzhen) (Approval number: CUHKSZ-AE2025006). The rats (n = 63) were randomly divided into seven experimental groups with 9 animals per group: control group, CTX group, CTX + melatonin (MT) group (MT is known to have ovarian protection effect (20) and used as the positive control), CTX + EGT group, CTX + low-dose product group, CTX + medium-dose product group and CTX + high-dose product group.

The CTX-induced POF rats model was established according to the previous studies (21). Animals were intraperitoneally injected with 50 mg/kg CTX on Day 1, followed by daily injections of 8 mg/kg CTX for 14 consecutive days. Rats in the control group received intraperitoneal injections of 0.9% saline instead of CTX during the same period. Rats in the control group and POF group received oral gavage of ultrapure water; the CTX + MT group received intraperitoneal injections of MT (30 mg/kg) dissolved in saline; the CTX + EGT group was treated with oral gavage of EGT (13 mg/kg) dissolved in ultrapure water. The CTX + low-, medium-, and high-dose product groups received the capsule components dissolved in ultrapure water by oral gavage at doses of 60 mg/kg, 120 mg/kg, and 240 mg/kg, respectively.

The low dose of the product (60 mg/kg) was calculated based on the equivalent recommended human daily intake (2 capsules each day). The weight of the active ingredients in each capsule is 285 mg (15 mg EGT, 41.5 mg CoQ10, 10 mg PQQ, 5.3 mg hydroxytyrosol, 100 mg vitamin E and 113.2 mg dextrin adjuvant). So the human dose is 570 mg for adults (60 kg body weight), which is 9.5 mg/kg. It is converted to rat dose 60 mg/kg using body surface area conversion. The medium and high doses were set at two-fold and four-fold of the low dose. The dose of EGT is selected based on its content in the high-dose product group. Because each capsule (285 mg) contains 15 mg EGT, which is 5.26%, EGT dose (13 mg/kg) is 5.26% of high dose product group (240 mg/kg). The dose of MT is selected based on previously reported studies (20).

The estrous cycle of rats was determined by performing vaginal smears, crystal violet staining, and microscopic observation daily for 15 consecutive days before the end of feeding (22). All rats were sacrificed under anesthesia with CO_2_. Then, blood, ovarian and skin were collected for further study.

### Biochemical analysis

Blood samples of rats were taken from inferior vena cava and then serum samples were obtained by centrifugation. The ELISA kits (Elabscience, Wuhan, China) were used to determine the levels of estradiol (E_2_), follicle stimulation hormone (FSH), luteinizing hormone (LH), anti-Müllerian hormone (AMH) and tumor necrosis factor Alpha (TNF-α), Interleukin 1 Beta (IL-1β) and Interleukin 6 (IL-6) according to the instructions of manufacturers. The measure of total superoxide dismutase (T-SOD), catalase (CAT) activity and malondialdehyde (MDA), glutathione (GSH) levels was performed using commercially available kits, T-SOD kit (A001-3), CAT kit (A007-1), MDA kit (A003-1) and GSH kit (A006-2) from Jiancheng Bioengineering Institute, Nanjing, China.

### Ovarian follicle counts and morphological analysis

Four ovarian samples from each group were randomly selected for histological analysis. Ovaries were fixed in 4% paraformaldehyde for 12 h, embedded in paraffin, and sectioned at a thickness of 3–5 μm continuously. Sections were mounted on glass slides and deparaffinized with xylene and stained with hematoxylin and eosin (H&E). Follicles were classified into primordial, primary, secondary, and atretic follicles according to the previously described criteria(23). Only follicles containing oocytes with clearly visible nuclei were included in the quantitative analysis of follicle numbers.

### TUNEL staining

Apoptosis of granulosa cells was determined using the In Situ Cell Death Detection Kit (Roche, USA) according to the manufacturer’s instructions. Following deparaffinization, sections were incubated with proteinase K (20 mg/mL) in a humidified chamber for 15 min and then treated with 3% H₂O₂ for 10 min to quench endogenous peroxidase activity. Subsequently, sections were incubated with TdT labeling buffer at 37 °C in a humidified chamber for 1 h, followed by counterstaining with DAPI. Five random fields per slide and four animals per group were selected for examination under a light microscope. The number of TUNEL-positive granulosa cells (stained brown) and the total number of granulosa cells (stained blue with DAPI) in antral follicles were counted in each field. The percentage of TUNEL-positive granulosa cells in antral follicles was quantified using Image-Pro Plus 6.0 software.

### Quantitative PCR (qPCR) Analysis

The total RNA of ten ovary samples of each group were extracted by RNAiso plus reagent (Takara Bio, Co. Ltd., Dalian, China) according to the manufacturer’s instructions. RNA concentration and purity were determined using a NanoPhotometer® N60 spectrophotometer (Implen, German). Subsequently, 1 μg of total RNA from each sample was reverse transcribed into cDNA using the PrimeScript™ RT reagent Kit (Takara, Japan). Real-time qPCR was performed using SYBR Green reagent (Thermo Scientific) on a QuantStudio 3 system (Applied Biosystems, USA). The primer sequences are provided in **Supplementary Table 1**. All reactions were performed in triplicate, and gene expression levels were normalized to GAPDH as an internal control.

### Statistical analysis

Data are presented as means ± standard deviation (SD). Data were analyzed by one-way ANOVA followed by Dunn’s post-test for multiple comparisons. GraphPad-Prism 9.0 (San Diego, CA, USA) was used to analyze data. Values of p < 0.05 were considered statistically significant.

## RESULTS

### The EGT and product could restore normal estrous cycle in CTX-treated rats

The estrous cycles of all rats were monitored daily by vaginal smears and microscopic examination. The average estrous cycle in the control group was 4 – 5 days. Compared with the control group, the cycle length was significantly extended in the POF model group after CTX treatment (**Figure 1A,1C**). EGT and the product groups in different doses significantly shortened the estrous cycle and restored the regular estrous cycle pattern compared with the POF model group (**Figure 1C**).

**Figure 1.**
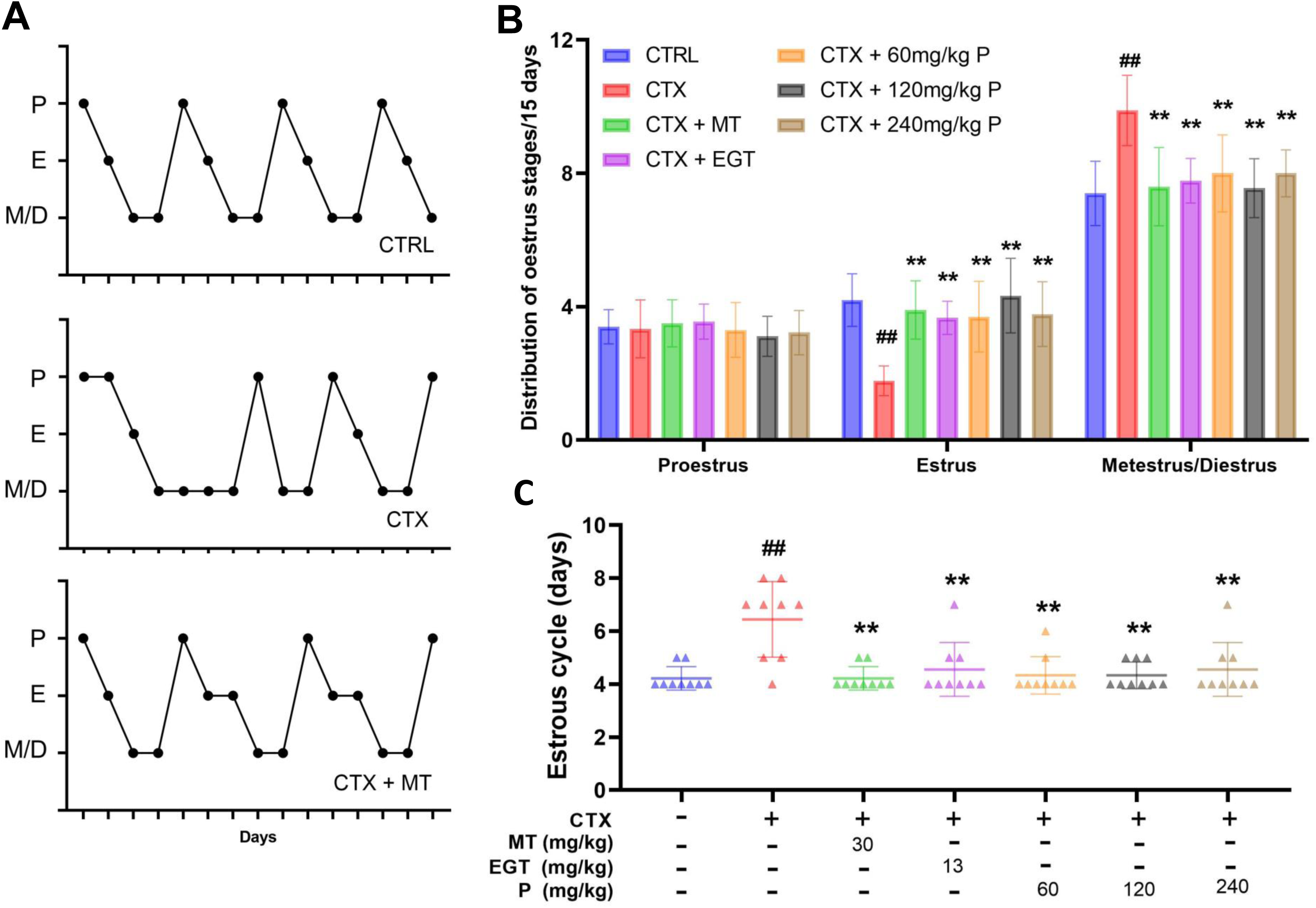
Changes in the estrous cycle of rats after 15 days of treatment. Proestrus (P), Estrus (E), Metestrus (M), and Diestrus (D) represent the four stages of the estrous cycle. (A) Representative estrous cycle patterns from the control group, the primary ovarian failure (POF) model group, and the treatment groups. (B) Distribution of time spent in each estrous stage over the 15-day period. (C) Length of the estrous cycle in different treatment groups. Data are presented as mean ± SD. ^##^P < 0.01 compared with the control group; *P < 0.05, **P < 0.01 compared with the POF group.

Furthermore, the number of days in proestrus, estrus, and metestrus/diestrus phases over a 15-day period was analyzed (**Figure 1B**). In the POF model group, the duration of estrus was significantly reduced, while the duration of metestrus/diestrus was markedly increased compared with the control group. The EGT and product groups in different doses significantly increased the number of estrus days (**Figure 1B**).

Ovarian weight was significantly decreased in the POF model group (CTX treatment group) compared with the control (**Figure 2D, 2E**). Notably, MET treatment, as well as the medium-and low-dose product, markedly reversed the CTX-induced decline in ovarian weight. EGT and low dose product increased ovary weight compared to CTX group but with no statistical significance. Co-administration of the product or EGT with CTX did not alter body weight or uterine weight compared with CTX alone.

**Figure 2.**
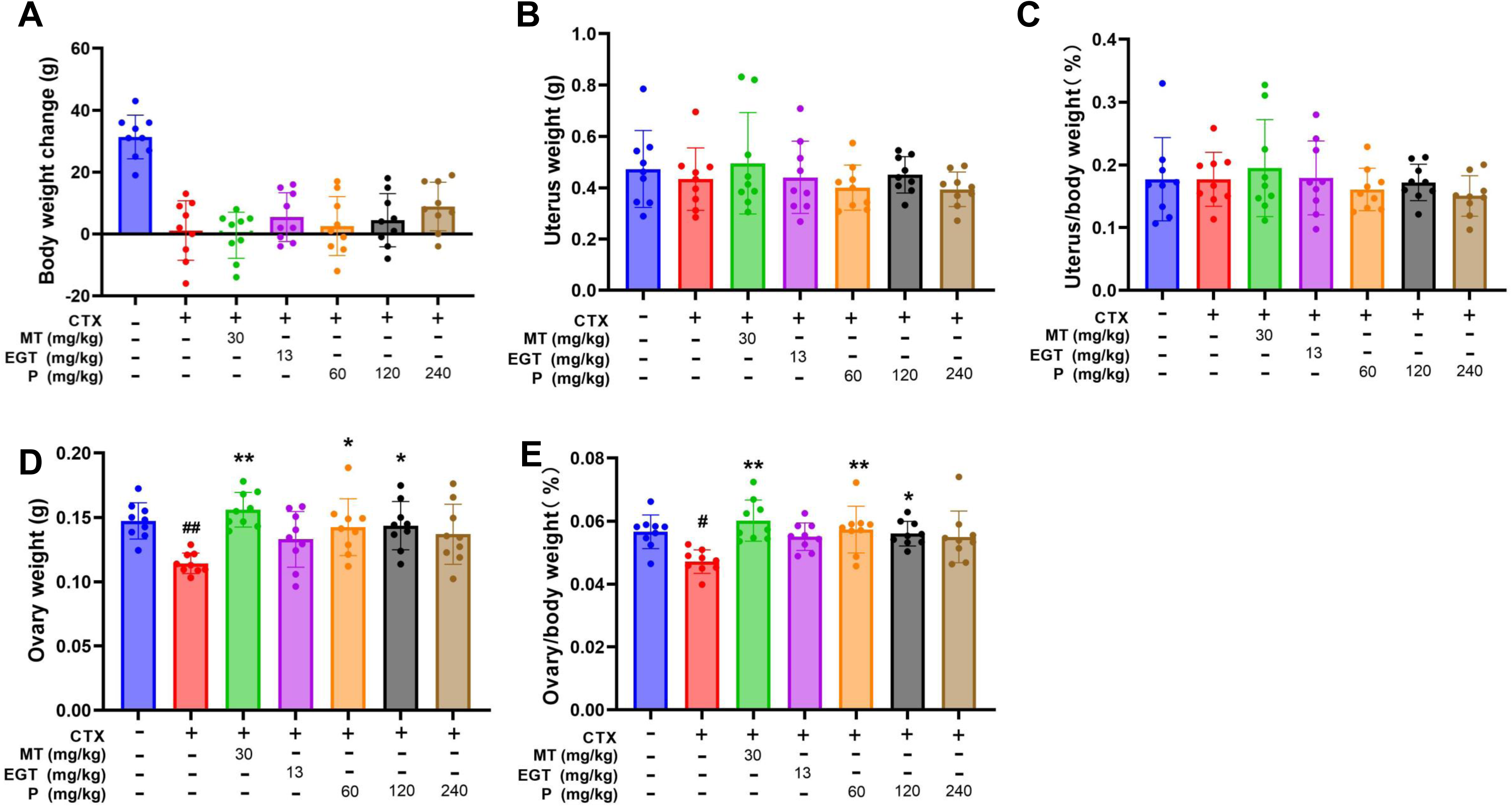
Changes in body weight, uterine weight, and ovarian weight in rats after 15 days of treatment. (A) Body weight of rats after 15 days of administration. (B) Uterine weight after 15 days of administration. (C) Uterine weight-to-body weight ratio. (D) Ovarian weight after 15 days of administration. (E) Ovarian weight-to-body weight ratio. Data are presented as mean ± SD. ^#^P < 0.05, ^##^P < 0.01 compared with the control group; *P < 0.05, **P < 0.01 compared with the POF group.

### The EGT and product restored normal ovary follicular development in rats

The numbers of follicles at different developmental stages (primordial, primary, secondary, and atretic) in each group were quantified using HE-stained slides (**Figure 3**). Compared with the control group, the CTX group exhibited a significant reduction in primordial follicle numbers (p < 0.01), decreasing to approximately 60% of the control, along with a significant increase in atretic follicles to nearly twice the control level (p < 0.05).

**Figure 3.**
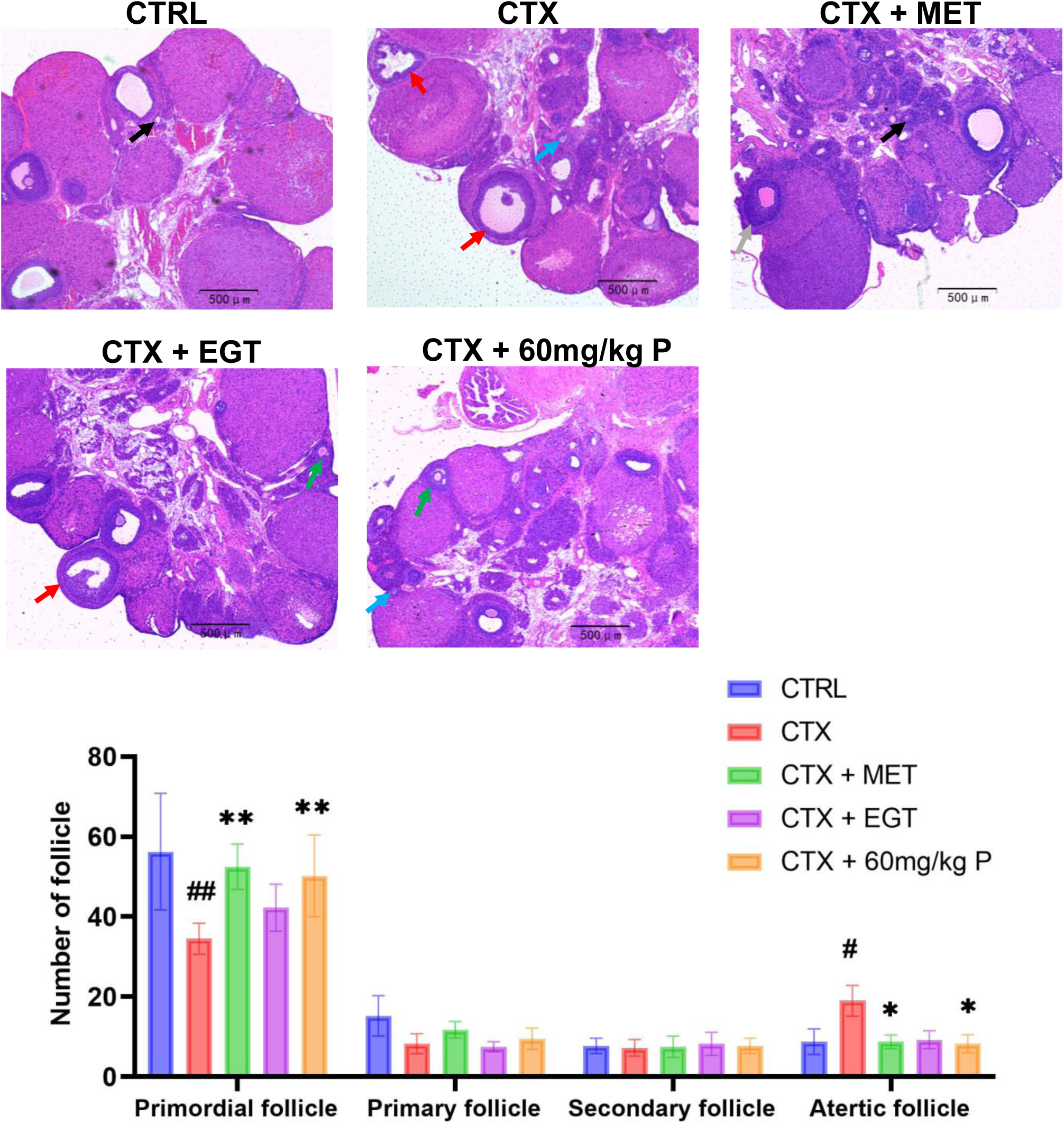
Effects of different interventions on follicular development in the POF model. Black arrows indicate primordial follicles, blue arrows indicate primary follicles, green arrows indicate secondary follicles, and red arrows indicate atretic follicles. Data are presented as mean ± SD.^#^P < 0.05, ^##^P < 0.01 compared with the control group; *P < 0.05, **P < 0.01 compared with the POF group.

Both MET and the low-dose product significantly increased the number of primordial follicles compared with the CTX group (p < 0.01), restoring them to normal levels comparable to those in the control group. Regarding atretic follicles, MET and low dose product significantly reduced the proportion of atretic follicles induced by CTX (p < 0.05), lowering them to control levels. EGT could also decrease the number of atretic follicles with no statistical significance.

To evaluate the effects of different interventions on granulosa cell apoptosis within atretic follicles, TUNEL staining was performed on ovarian sections from each group. Compared with the control group, the CTX group exhibited a significantly higher percentage of TUNEL-positive cells (p < 0.01), indicating that the CTX-induced POF model markedly induced granulosa cell apoptosis (**Figure 4**). Treatment with MET, EGT, or the low-dose product significantly reduced the proportion of TUNEL-positive cells compared with the CTX group (p < 0.01), demonstrating that all they effectively reduced CTX-induced granulosa cell apoptosis.

**Figure 4.**
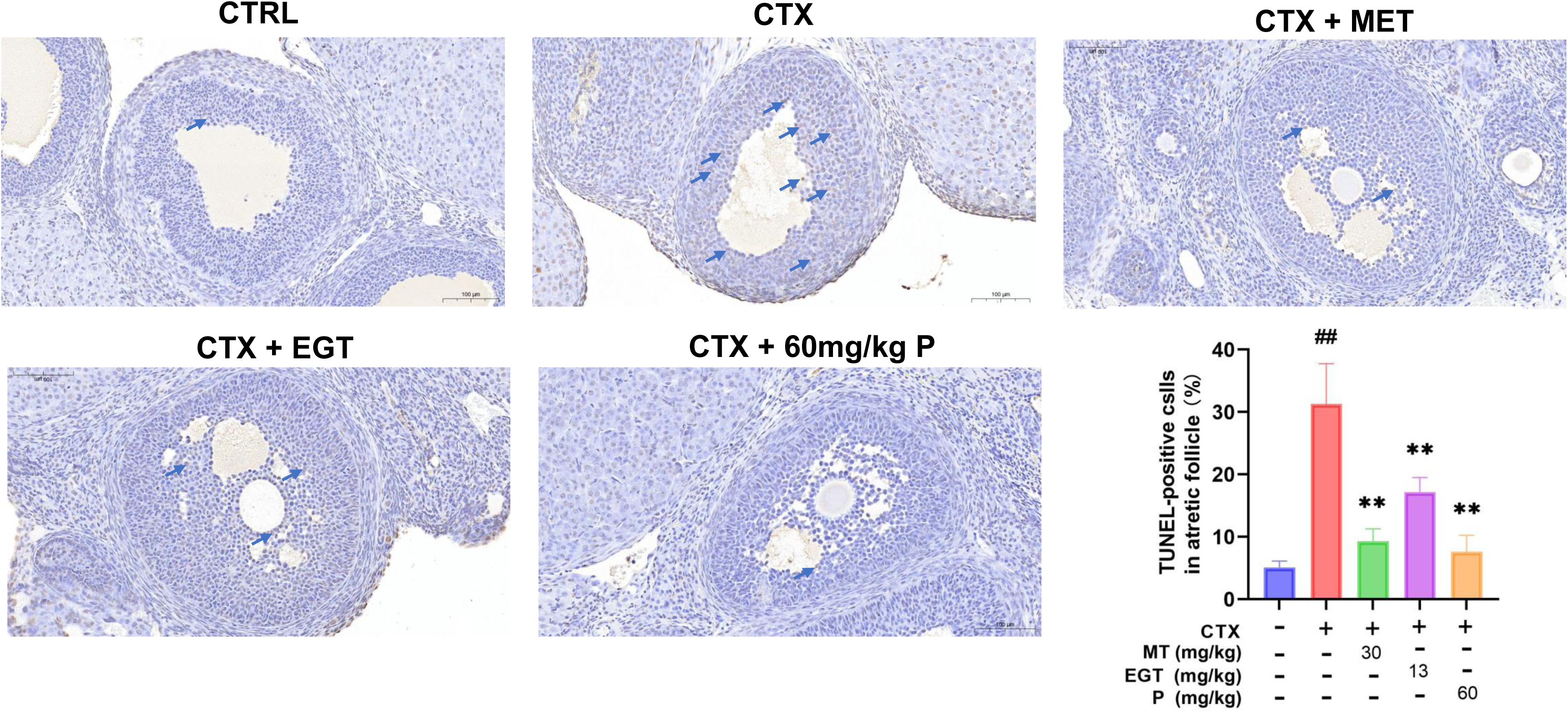
Percentage of TUNEL-positive cells in atretic follicles of rat ovaries after 15 days of treatment. Apoptotic cells were assessed by in situ TUNEL assay. In this assay, fragmented DNA in ovarian cells was labeled with horseradish peroxidase and visualized by color development using the DAB substrate. TUNEL-positive apoptotic nuclei appear brownish-yellow (indicated by blue arrows). Data are presented as mean ± SD. ^##^P < 0.01 compared with the control group; P < 0.01 compared with the POF group.

### The EGT and product restored serum hormone levels in CTX treated rats

To further assess the effects of EGT and product on ovarian function, serum levels of multiple hormones related to ovarian physiology were measured, including estradiol (E₂), anti-Müllerian hormone (AMH), luteinizing hormone (LH), and follicle-stimulating hormone (FSH). E₂ and AMH levels were significantly decreased in the CTX-treated group compared with the control (p < 0.01), whereas LH and FSH levels were markedly elevated (p < 0.01) (**Figure 5**), indicating impaired ovarian reserve and dysregulation of the hypothalamic–pituitary–ovarian (HPO) axis.

**Figure 5.**
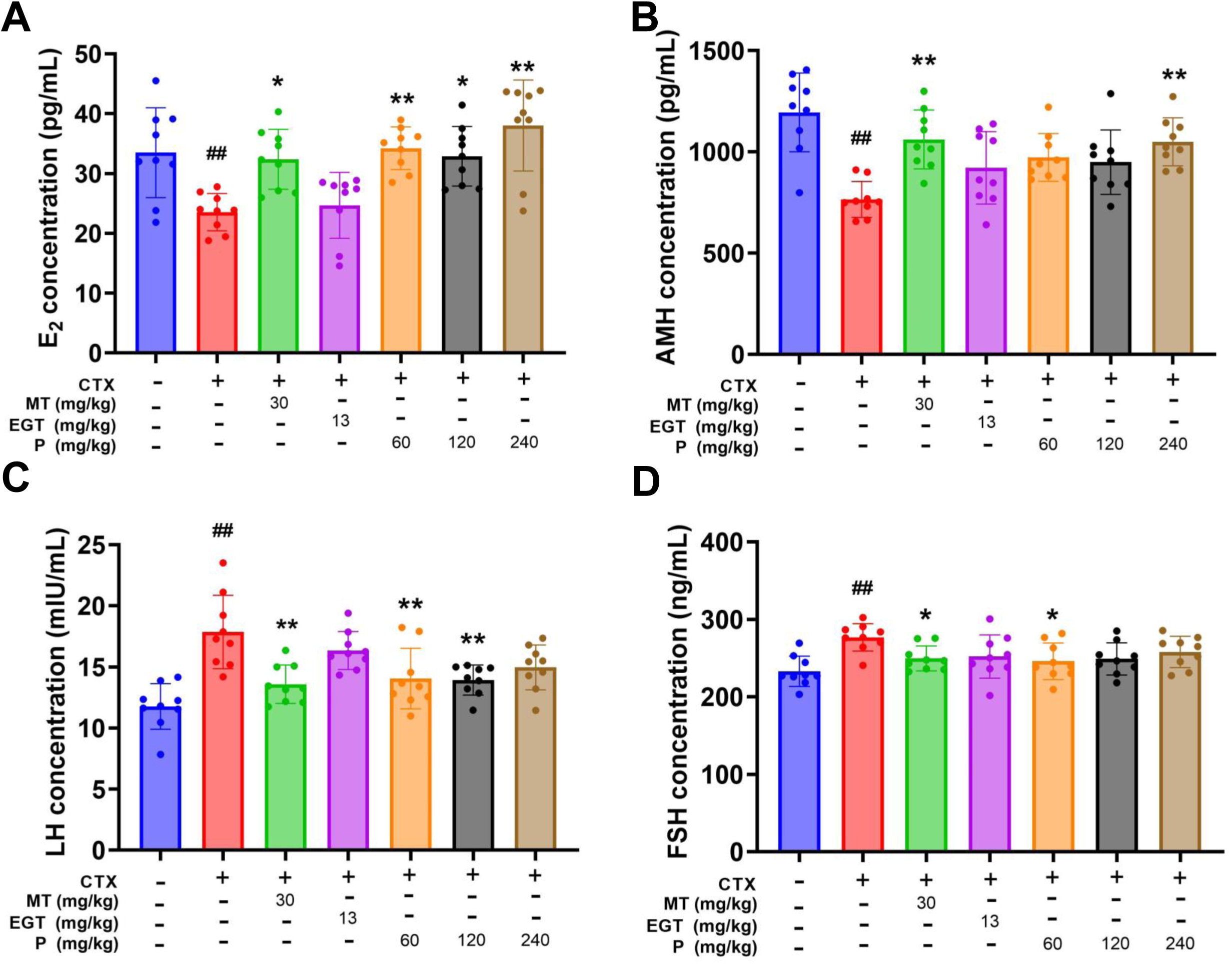
Effects of different interventions on serum sex hormone levels in POF model rats. (A) Estradiol (E₂), (B) anti-Müllerian hormone (AMH), (C) luteinizing hormone (LH) and (D) follicle-stimulating hormone (FSH) concentrations. Data are presented as mean ± SD. ^##^P < 0.01 compared with the control group; *P < 0.05, P < 0.01 compared with the POF group.

Treatment with the positive control MET significantly increased serum E₂ and AMH levels (**Figure 5A, 5B**) while markedly reduced the elevated LH and FSH levels (**Figure 5C, 5D**) in POF rats. Similarly, administration of the product at low, medium, or high dosages elevated E₂ and AMH levels and decreased LH and FSH levels compared with the CTX group.

### The EGT and product decreased oxidative stress levels in CTX treated rats

Several representative oxidative stress-related parameters in ovary were measured, including catalase (CAT), superoxide dismutase (SOD) activity, glutathione (GSH), and malondialdehyde (MDA). CAT activity, SOD activity and GSH level in ovary was significantly reduced following CTX induction, indicating impaired antioxidant capacity (**Figure 6A-6C**). The product treatment significantly increased CAT activity, SOD activity and GSH levels (**Figure 6A-6C**). EGT could also restore CAT activity, SOD activity and GSH levels to some extent compared to CTX alone group with no statistical significance.

**Figure 6.**
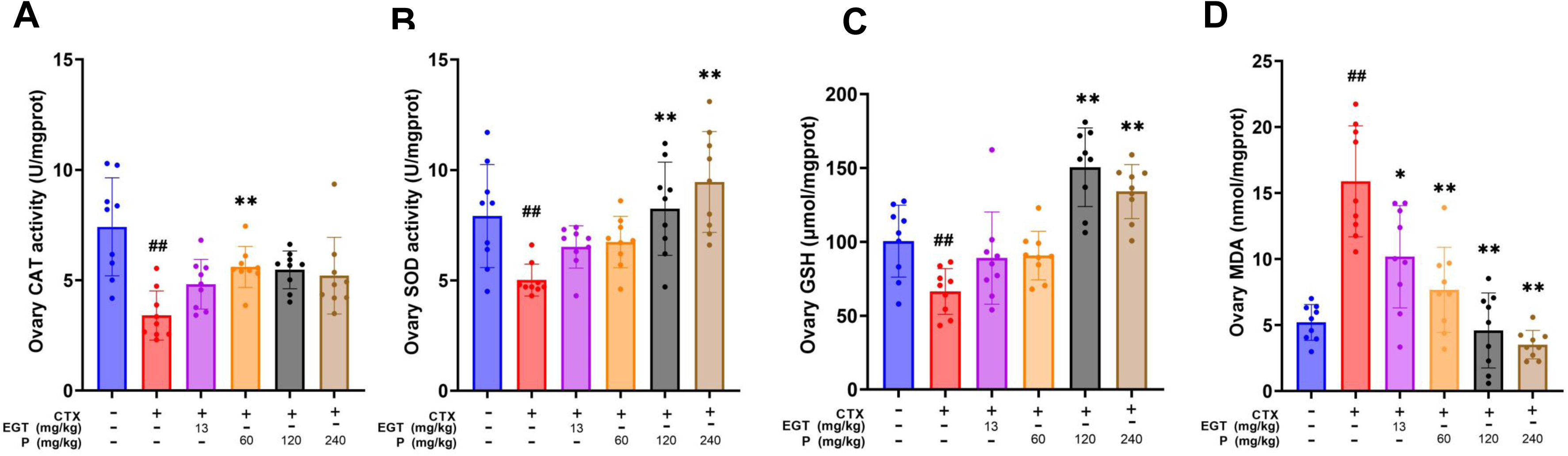
Effects of different interventions on ovarian oxidative stress markers in POF model rats. (A) catalase (CAT) activity, (B) superoxide dismutase (SOD) activity, (C) glutathione (GSH) level, (C) malondialdehyde (MDA) level, (D). Data are presented as mean ± SD. ^##^P < 0.01 compared with the control group; *P < 0.05, P < 0.01 compared with the POF group.

CTX treatment significantly increased ovarian MDA content compared with the control group, indicating a marked elevation in lipid peroxidation (**Figure 6D**). Compared with the CTX group, EGT and all doses of the product significantly reduced MDA concentrations, with the medium-and high-dose product groups showing reductions approaching or below control levels (**Figure 6D**).

To investigate the potential regulatory effects of different interventions on ovarian mitochondrial function, we examined the mRNA expression levels of mitochondrial biogenesis–related genes *OPA1*, *Mfn1*, and *Mfn2* (24) in ovarian tissues from each group. CTX treatment did not cause significant changes in the expression of these three genes compared with the control group (p > 0.05), suggesting that CTX did not markedly suppress the basal expression of mitochondrial biogenesis–related genes in the ovary (**Figure 7**). However, rats receiving the low-dose product exhibited significantly higher expression levels of *OPA1*, *Mfn1*, and *Mfn2* compared with the CTX group (**Figure 7**), indicating that the low-dose product enhanced the activity of mitochondrial biogenesis–related pathways. No significant changes were observed in the expression of these genes in the EGT group, indicating that the effect of product in these gene expression is due to the other components in the product other than EGT.

**Figure 7.**
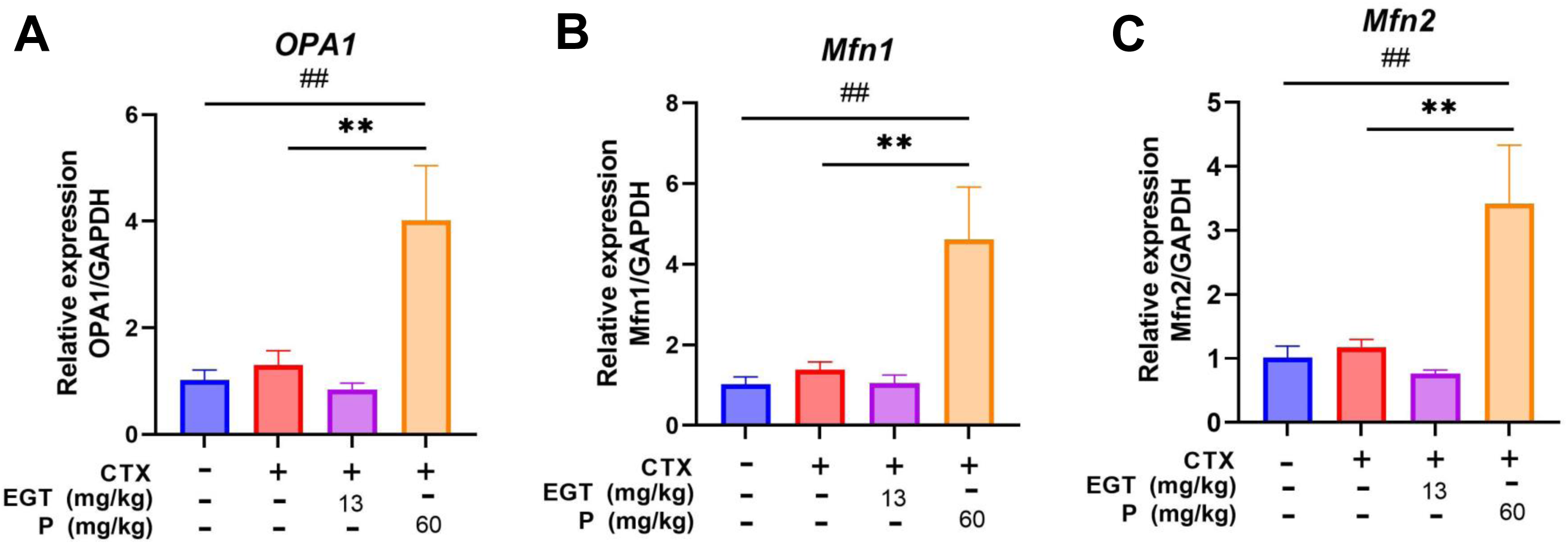
Expression levels of mitochondrial biogenesis-related genes in the ovaries of rats after different interventions. (A) *OPA1*, (B) *Mfn1*, and (C) *Mfn2* expression in ovarian tissue. Data are presented as mean ± SD. ##P < 0.01 compared with the control group; P < 0.01 compared with the POF group.

### EGT and product could ameliorate skin inflammation induced by CTX in rats

To investigate the impact of CTX-induced POF on local skin inflammatory status and to evaluate the anti-inflammatory effects of the interventions, the levels of tumor necrosis factor-α (TNF-α), interleukin-6 (IL-6), and interleukin-1β (IL-1β) in skin tissues from each group were measured (**Figure 8**). TNF-α levels were significantly higher in the CTX group compared with the control group, indicating that ovarian dysfunction may trigger systemic or local pro-inflammatory responses. Both EGT and the low-dose product significantly reduced TNF-α levels relative to the CTX group, suggesting a marked inhibitory effect of EGT and the low-dose product on TNF-α expression. EGT treatment significantly decreased IL-6 and IL-1 β levels compared to CTX group (**Figure 8B, 8C**), while the low-dose product also decreased IL-6 and IL-1 β levels compared to CTX group although with no statistical significance.

**Figure 8.**
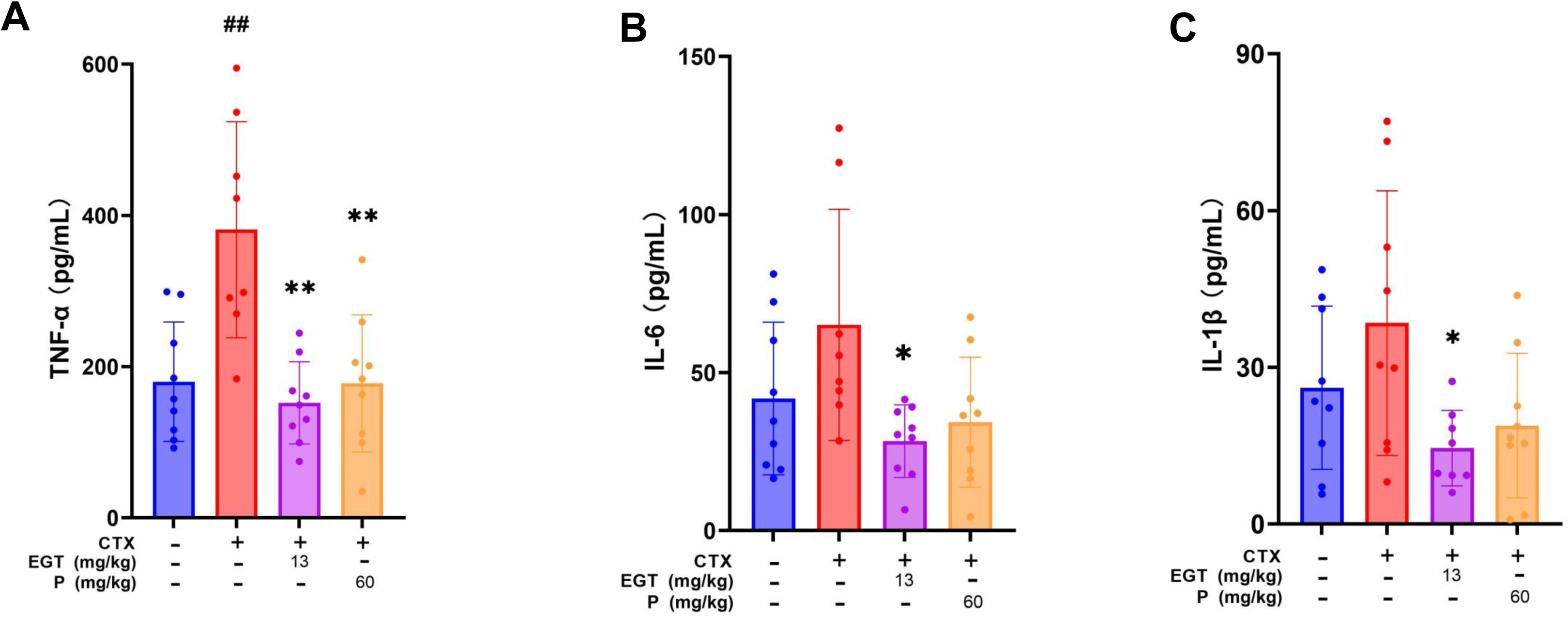
Changes in inflammatory cytokine levels in rat skin tissue after 15 days of treatment. (A) Tumor necrosis factor-α (TNF-α), (B) interleukin-6 (IL-6), and (C) interleukin-1β (IL-1β) concentrations in skin tissue. Data are presented as mean ± SD. ##P < 0.01 compared with the control group; *P < 0.05, P < 0.01 compared with the POF group.

Skin collagen fiber morphology was examined by Masson’s trichrome staining, and the collagen-to-muscle fiber ratio (C/M ratio) was quantified to assess the effects of different treatments on skin structure (**Supplementary Figure 1**). In the control group, the C/M ratio was the highest, with abundant blue-stained collagen fibers evenly distributed throughout the dermis. The CTX group showed a marked reduction in the C/M ratio and decreased collagen fiber density, although without statistical significance. EGT and the low-dose product increased the C/M ratio to some extent relative to the CTX group (**Supplementary Figure 1**), suggesting that both interventions provide partial protection of dermal collagen structure.

## DISCUSSION

This study evaluated the therapeutic effects of the product and one of its active constituents, EGT, on CTX-induced POF in a rat model. We found that the product and EGT could restore the normal estrous cycle, ovarian morphology and function, hormonal balance, antioxidant status and inhibit skin-associated inflammation.

CTX markedly prolonged the estrous cycle, reduced ovarian weight, accompanied by decreased serum E₂ and AMH levels, elevated LH and FSH levels—hallmarks of diminished ovarian reserve and gonadal axis dysregulation (25, 26), suggesting the disruption of hypothalamic–pituitary–ovarian axis function, consistent with previous reports (22, 27). The product and EGT significantly shortened the cycle, restored the duration of the estrus phase, increased E₂ and AMH concentrations while reduced LH and FSH levels, implying effectiveness in improving pituitary–ovarian axis activity, and preserving ovarian endocrine function.

The initial pool of primordial follicles is established at birth and is non-renewable, serving as the finite reservoir that supports a woman’s entire reproductive lifespan (28).

During folliculogenesis, oocytes undergo a series of developmental stages (primordial, primary, secondary, and mature) culminating in ovulation and the potential for reproduction (29). However, the vast majority of primordial follicles undergo atresia, and only a small subset is destined to mature (30). Histomorphological evaluation showed that CTX significantly reduced the number of primordial follicles and increased the proportion of atretic follicles, reflecting depletion of follicular reserves and accelerated apoptosis. The product (particularly effective at low doses) and EGT treatment increased the number of primordial follicles and reduced follicular granulosa cell apoptosis, indicating their protective effects on follicle development. AMH serves as a critical early biomarker of ovarian aging, and its levels progressively decline during the course of ovarian senescence in rats (31). The parallel increase in AMH levels further supports the role of the product and EGT in maintaining ovarian function.

In this study, CTX administration led to significant increases in serum FSH and LH levels (**Figure 5C, 5D**), a marked reduction in serum estradiol (E_2_) levels (**Figure 5A**) compared with the control group. Notably, these hormonal changes were effectively rescued by oral administration of the product at various doses. These findings suggest that the product is effective in restoring ovarian dysfunction and the hypothalamic–pituitary–gonadal (HPG) axis.

Oxidative stress is a pivotal contributor to POF pathogenesis (32,33). In the CTX model, activities of key ovarian antioxidant enzymes (CAT and SOD) were significantly reduced, while MDA, a lipid peroxidation product, was elevated, indicating oxidative imbalance. Product and EGT intervention restored CAT activity, GSH content, and SOD levels, while reducing MDA, with the most pronounced effects observed in the medium-and high-dose groups. These results suggest potent antioxidative properties, which may represent one possible mechanism by which the product and EGT improve ovarian function.

In addition to reproductive impairment, ovarian dysfunction is closely associated with skin aging (34). In addition, as reported, CTX could cause skin damage (35). In this study, CTX elevated inflammatory cytokines (TNF-α, IL-6, IL-1β) in skin tissue. Nevertheless, the product and EGT intervention reduced TNF-α and IL-1β levels, revealing anti-inflammatory potential, with implications of the product and EGT for applications in skin health maintenance.

In conclusion, this study elucidates the possible mechanisms by which the product and EGT mitigate CTX-induced POF, including hormonal regulation, oxidative stress alleviation, follicular preservation, and local anti-inflammatory activity in skin. The bioactivity observed in the low-dose intervention group highlights its promising potential for ovarian function preservation and delaying ovarian aging. Further investigations are merited to illustrate the detailed mechanism for EGT’s protective effects in POF.

---

**Supplementary Figure 1.**
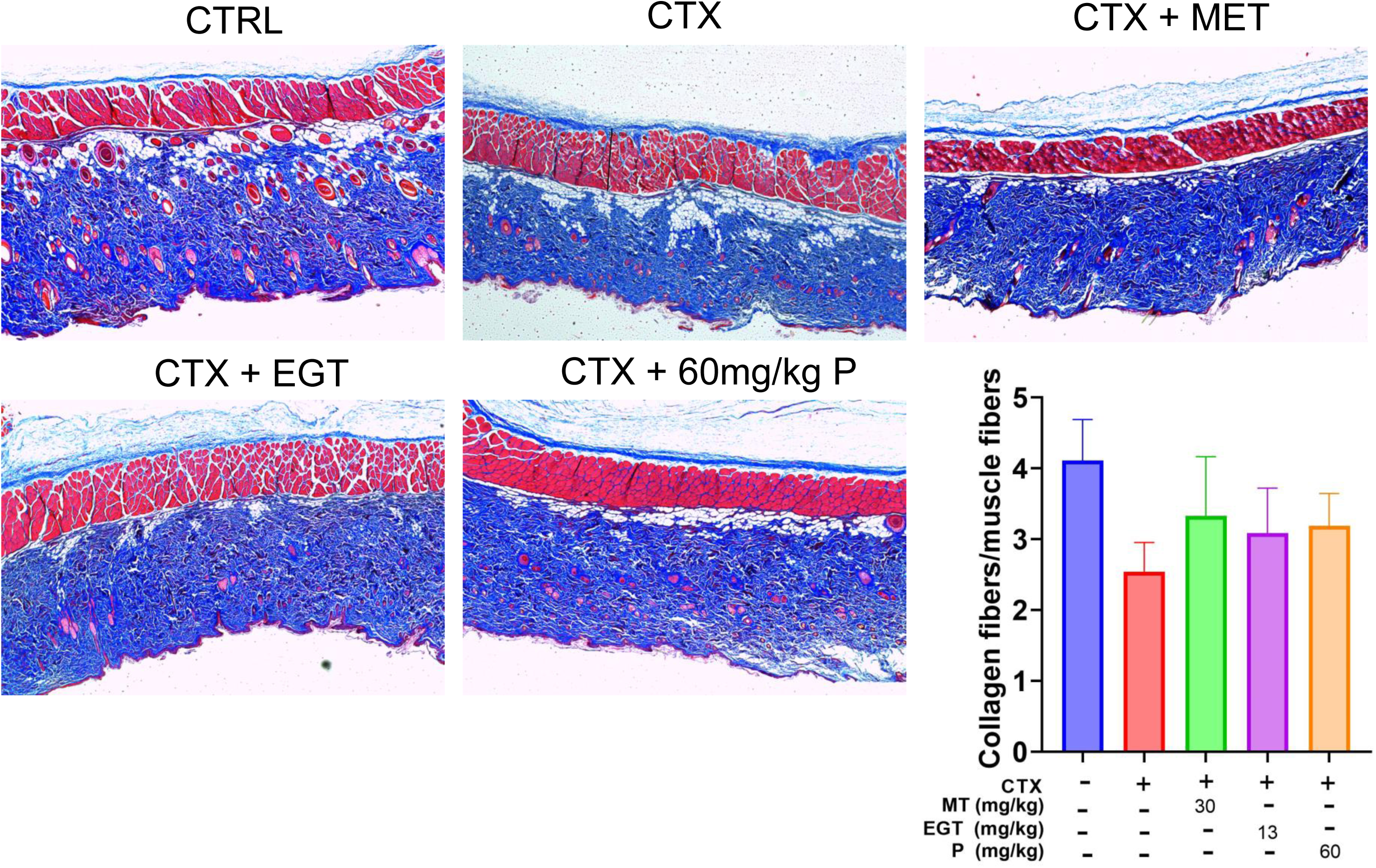
Changes in the collagen-to-muscle fiber ratio (C/M ratio) in rat skin tissues after 15 days of treatment. Masson’s trichrome staining shows muscle fibers in red and collagen fibers in blue. Data are presented as mean ± SD.

**Supplementary Table 1.**
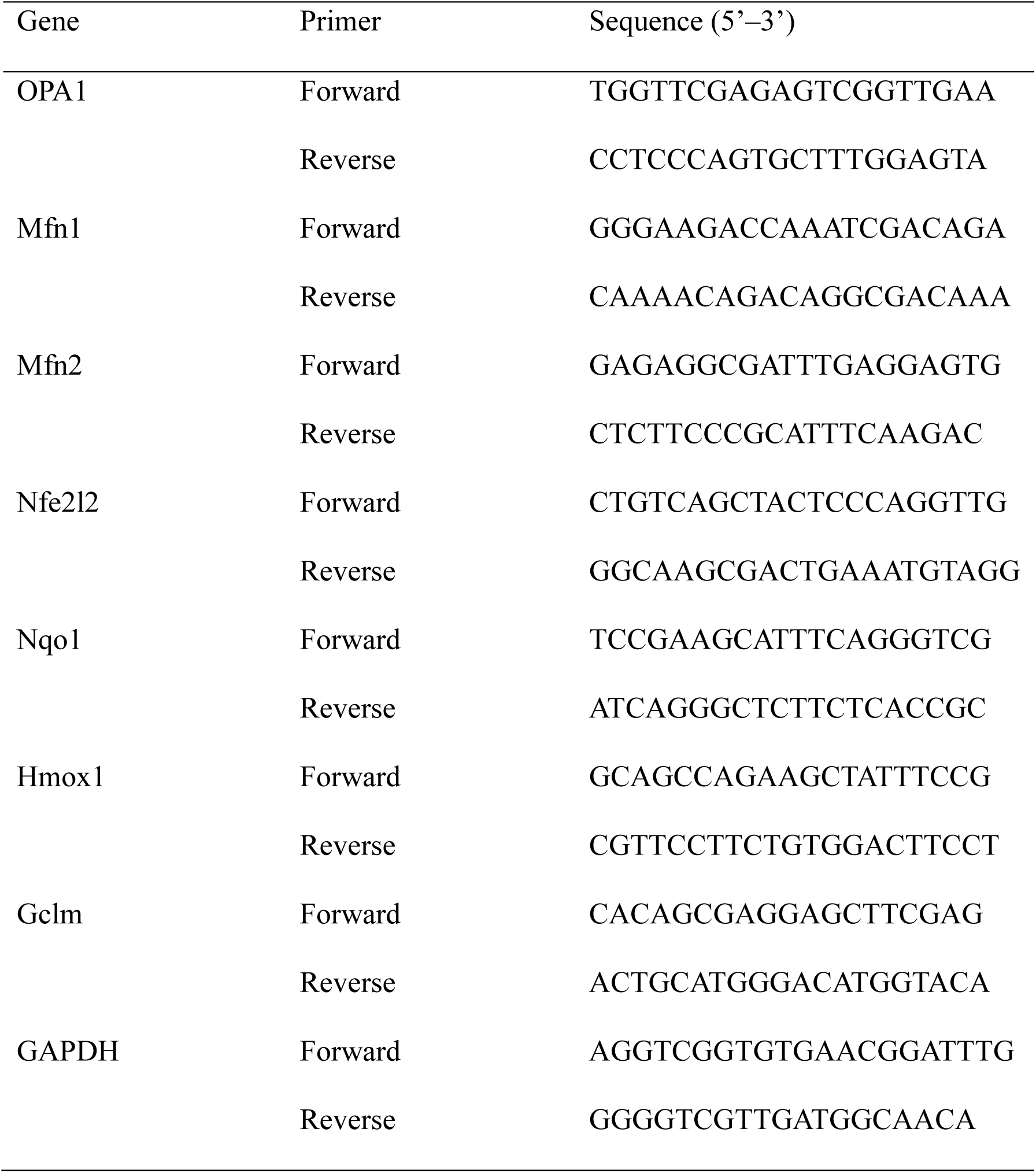
Genes and primers used in the RT-qPCR.

